# Myeloid cell recruitment propels right ventricular dysfunction in HFpEF via sterile inflammation

**DOI:** 10.1101/2025.11.21.689863

**Authors:** Lara Jaeschke, Ceren Koçana, Alexandra Maria Chitroceanu, Annika Winkler, Hannah Kleitke, Pauline Fahjen, David Faidel, Erik Asmus, Martin Meiser, Efstathios G Stamatiades, Karolin W. Hublitz, Veronika Zach, Lucie Kretzler, Daniela Zurkan, Paul-Lennard Perret, Leonardo A. von der Ohe, Szandor Simmons, Jonathan L. Gillan, Kristina Franz, Lina Alasfar, Virginia S. Hahn, Kavita Sharma, Edwardo Reynolds, Gabriele G. Schiattarella, Sophie van Linthout, Burkert Pieske, Niklas Hegemann, Philipp Mertins, Frank Edelmann, Wolfgang M. Kuebler, Jana Grune

## Abstract

**Background:** In contrast to what has already been shown in HFpEF associated left ventricular (LV) diastolic dysfunction, leukocytes’ role in frequently occurring right ventricular dysfunction (RVD) secondary to HFpEF are so far missing, partially due to the lack of suitable small animal models. Here, we follow a translational research approach by establishing a murine HFpEF model developing manifest RVD and analyzed human HFpEF cohorts to study the mechanistic link between leukocytes and RVD in HFpEF.

**Methods:** 8-week-old male and female *C57BL/6J* or *Cx3cr1^CreER^/+R26^tdTomato/+^* mice were divided into four experimental groups: i) chow, ii) HFpEF (N[ω]-nitro-l-arginine methyl ester (L-NAME), 60% high-fat diet), iii) chronic hypoxia (10% O_2_) and iv) HFpEF and hypoxia (RV-HFpEF) to assess bi-ventricular function and myeloid cell dynamics. To test whether myeloid cells are causally involved in the development of RV remodeling in HFpEF, we additionally treated RV-HFpEF mice with the colony stimulating factor 1 receptor inhibitor PLX-5622 (PLX) to deplete myeloid cells. After 12 weeks, all experimental groups were subjected to transthoracic echocardiography, invasive hemodynamics or flow cytometry.

**Results:** RV-HFpEF resulted in LV diastolic dysfunction indicated by increased E/E’ ratio, reduced global longitudinal peak strain, smaller end-diastolic diameters and increased isovolumetric relaxation time compared to chow. RV-HFpEF animals developed RV hypertrophy and RVD evident as increased Fulton’s index and collagen content as well as elevated RV systolic pressures (RVSPs) and reduced tricuspid annular plane systolic excursion, respectively. Flow cytometric analyses revealed elevated total leukocyte, monocyte, and macrophage counts in RV tissue of RV-HFpEF compared to chow or LV tissue from RV-HFpEF animals. These data were confirmed by unbiased proteomic analyses of RV tissue from RV-HFpEF mice, demonstrating increased abundance of proteins involved in activation of the innate immune system, macrophage chemotaxis, cell adhesion and extracellular matrix organization when compared to LV tissue or other experimental groups. Fate mapping experiments revealed that recruited monocyte-derived macrophages became the main source of total cardiac macrophages in RV tissue from RV-HFpEF mice. Depletion of myeloid cells was associated with rescued RVSP profiles compared to RV-HFpEF control mice. In HFpEF patients, RV dilation was associated with an increased percentage of circulating monocytes. In RV biopsies from HFpEF patients, we found increased expression of adhesion molecules, fibrotic markers and inflammatory transcripts.

**Conclusion:** We demonstrate that dysregulated myeloid cell dynamics are associated with, and directly contribute to, the pathogenesis of HFpEF-associated RVD in humans and mice.

**Clinical Perspective:** **What is new:**

- We explore myeloid cell dynamics in a novel three-hit experimental HFpEF mouse model with RV hypertrophy, RV end-systolic pressure and RV dysfunction.
- In this model, RV dysfunction was associated with macrophage expansion, monocyte recruitment and extracellular matrix deposition, whilst macrophage depletion partly reversed these changes and rescued RV hemodynamics.
- HFpEF patients with RV dilation or RV dysfunction exhibit unique leukocyte dynamics and inflammatory profiles when compared to HFpEF patients with normal RV function or diameters.

**Clinical implications:**

- There exists a major clinical discrepancy between high incidence of RV dysfunction associated to HFpEF and a lack of targeted treatment strategies.
- Our novel three-hit mouse model recapitulates many features of the clinical scenario of HFpEF patients with RV dysfunction, therefore representing an important step towards systematic testing and development of targeted treatment options.
- Sterile inflammation and dysregulation of innate immune cells may be suitable targets for therapeutic interventions against RV dysfunction in HFpEF.

## Introduction

Heart failure with preserved ejection fraction (HFpEF) is frequently associated with risk factors and comorbidities such as hypertension, higher age, sedentary lifestyle, and obesity^1,2^. The synergism of hemodynamic and metabolic stress in concert with low-grade chronic inflammation is considered to contribute to disease manifestation and progression in the left ventricle (LV)^3,4^. Consistently, proinflammatory markers such as interleukin-6, tumor necrosis factor-alpha or high-sensitivity C-reactive protein have been established as predictors of clinical outcome in HFpEF patients, pointing towards a causal role of inflammation in HFpEF pathogenesis^5^. As of yet, macrophage expansion sourced by increased hematopoiesis and myeloid cell recruitment in HFpEF LV tissue, is widely accepted as a pathophysiological hallmark of diastolic dysfunction^6,7^. Of note, this HFpEF-associated dysregulation of systemic, sterile inflammation does not exclusively affect the LV, but also tissues and organs beyond the LV, such as the pulmonary microvascular system, the gut microbiome, and the kidneys^8–10^.

HFpEF patients frequently develop secondary pulmonary hypertension and subsequent right ventricular dysfunction (RVD)^11–13^. Importantly, RVD is recognized as a negative prognostic factor in LV failure, independent of the underlying cause^14^. This ventricular interdependence has traditionally been attributed to increased RV afterload secondary to pulmonary congestion, leading to RV remodeling, dilation and dysfunction, resulting in shortness of breath, fatigue, and syncope in affected patients^15^. This paradigm has, however, been challenged by the fact that RVD is primarily related to LV dysfunction and not to systolic pulmonary artery pressure *per se*. Moreover, RVD has recently been observed in HFpEF patients in the absence of PH^16^, suggesting that chronic pulmonary artery pressure elevation alone is insufficient to explain RVD in HFpEF^17,18^. As such, mechanisms other than direct hemodynamic forces, such as interventricular signaling, metabolic effects, or inflammatory cues have been proposed to contribute to RVD in LV failure^19,20^, but little is known about the underlying pathophysiological mechanisms fueling the development of RVD in HFpEF.

Here, we investigate the role of myeloid cells in RVD in HFpEF, following the notion that the innate immune system and dysregulated sterile inflammation are key pathogenic drivers of HFpEF-associated RVD. Since small animal models fully recapitulating the clinical HFpEF syndrome are rare and often poorly phenotyped for RV function and dimensions^21^, we employed a novel mouse model of concomitant hypertensive, metabolic and hypoxic stress phenocopying RVD development in HFpEF. In this model, myeloid cell accumulation, proliferation and recruitment were associated with, and directly contributed to, the pathogenesis of RVD in HFpEF mice via RV remodeling and tissue inflammation. Moreover, we show that HFpEF-RVs exhibit unique pathogenic responses compared to HFpEF-LVs, when exposed to the same systemic stressors, with HFpEF-RVs being particularly vulnerable to myeloid cell recruitment and extracellular matrix deposition. In the clinical scenario, we found that HFpEF patients with RV dilation had unique white blood cell signatures with a higher percentage of blood monocytes when compared to HFpEF patients with regular RV diameters. RV biopsies from HFpEF patients exhibited inflammatory and pro-fibrotic signatures. Viewed together our data indicate a causal role for myeloid cells in HFpEF-associated RVD.

## METHODS

Detailed methods are provided in the online supplement.

### Human

#### HFpEF definition

HFpEF was defined in accordance with current ACC diagnostic guidelines^22^ as the presence of the following symptoms and signs: i) at a minimum New York Heart Association (NYHA) Class II, ii) a LVEF greater than 50%, and iii) objective evidence of cardiac structural abnormalities [left ventricular mass index (LVMI) > 95 g/m² in females and > 115 g/m² in males, relative wall thickness (RWT) > 0.42, left atrial volume index (LAVI) > 34 mL/m²] and/or functional abnormalities [E/e’ ratio at rest > 13, pulmonary artery systolic pressure (PASP) > 35 mmHg] indicative of left ventricular diastolic dysfunction or elevated left ventricular filling pressures, and iv) elevated levels of natriuretic peptides [N-terminal pro-B-type natriuretic peptide (NT-proBNP) > 125 pg/mL].

#### CRC1470 cohort

HFpEF patients were prospectively enrolled in the clinical cohort of the Collaborative Research Center 1470 (CRC1470), a single-center consortium funded by the German Research Foundation (DFG: Deutsche Forschungsgemeinschaft). The study was conducted in accordance with the Declaration of Helsinki and approved by the Charité – Universitätsmedizin Berlin (Berlin, Germany) Ethics Committee (ethical approval EA1/224/21). All participants were over 18 years old and provided written informed consent. Patients were excluded if they had a life expectancy of less than one year, a history of heart failure with mid-range or reduced ejection fraction, acute coronary syndrome within the last 30 days, cardiac surgery within the past three months, or end-stage renal disease with or without hemodialysis. Additional criteria encompassed severe valvular heart disease, hypertrophic cardiomyopathy, amyloidosis, congenital heart diseases, sarcoidosis, and constrictive pericarditis. Patients with extracardiac conditions that could explain their symptoms, such as chronic obstructive pulmonary disease (COPD) GOLD stage > 2, primary pulmonary hypertension, or moderate to severe anemia (hemoglobin < 10 g/dL in males and < 9.5 g/dL in females) were also excluded. All patients underwent a comprehensive clinical evaluation, which included a two-dimensional echocardiography (2DE).

#### Bulk RNA-sequencing of human HFpEF biopsies

All data were pulled from a previously published data repository^23^. In brief, human HFpEF samples were obtained from patients who underwent endomyocardial biopsy of the RV septum at Johns Hopkins University. HFpEF diagnosis was based on consensus criteria, as described previously^23^. Control RV septal tissue came from unused donor hearts from the University of Pennsylvania. Differential expression testing was performed with DESeq2 as previously described^23^ with removal of the filter for low expression. The normalized expression of selected genes was graphically illustrated.

### Mice

All animal studies performed are conform to Directive 2010/63/EU of the European Parliament, the guidelines of the German Law on the Protection of Animals and the Guide for the Care and Use of Laboratory Animals published by the US National Institutes of Health (NIH Publication No. 85-23, revised 1985). The experimental protocols were reviewed and approved by the local authorities (Landesamt für Gesundheit und Soziales, Berlin, Germany; protocol no. G0008/22 and G0025/24) and performed in agreement with the Animal Research: Reporting of In Vivo Experiments (ARRIVE) guidelines. Wildtype mice were purchased from Janvier-Labs. *Cx3cr1^CreER/+^R26^tdTomato/+^* mice were kindly provided by Efstathios Stamatiades (Charité Berlin) for fate-mapping experiments and were generated by crossing *Cx3cr1^CreER/+^* with *R26^tdTomato/tdTomato^* mice^24,25^.

#### RVD induction in HFpEF mice

Mice were housed for at least 1 week to acclimatize to laboratory conditions before starting experimental procedures. We housed two to four mice per cage, divided by sex and treatment, kept in a 12-h:12-h light/dark cycle with free access to food and water in individually-ventilated cages and specific pathogen-free rooms. All mice underwent body weight monitoring twice a week and daily health monitoring.

Eight week-old male and female *C57BL/6J* mice were divided into four experimental groups: i) naive mice receiving standard diet (chow, 9% fat, Ssniff) and regular drinking water, serving as controls, ii) mice receiving a rodent diet with 60% fat (high-fat diet, Ssniff) and Nω-nitro-L-arginine methylester hydrochloride (L-NAME, 0.5 g/L, N5751-10G, Sigma-Aldrich) diluted in drinking water for twelve weeks to induce HFpEF, iii) mice receiving high-fat diet, L-NAME diluted in drinking water with additional exposure to chronic hypoxia (10% O_2,_ RV-HFpEF) for two weeks starting from week 10 to induce RVD in HFpEF, iv) chow mice exposed to hypoxia for two weeks were age- and sex-matched and were run in parallel with all other experimental groups.

### Statistics

Statistical analyses for human cohorts and proteomics were performed using R or SPSS. For two-group comparisons, we used Wilcoxon rank sum test. For categorical variables, we used Pearson Chi-squared test or Fisher’s exact test. Proteomic spectra were processed with Spectronaut, and quantified protein groups were filtered for completeness and quality before normalization and imputation of missing values. Differential expression was assessed with limma using duplicate correlation, and pathway enrichment was evaluated by single-sample GSEA (ssGSEA) based on the Reactome pathway database with Benjamini–Hochberg correction for multiple testing. For mouse studies, statistical analyses were performed using GraphPad Prism 9. Results are reported as box plots using the 25th (Q1), 50th (median or Q2) and 75th (Q3) percentiles and the interquartile range (IQR = Q3 − Q1), which covers the central 50% of the data. Sample sizes below 20 were assumed to be not normally distributed by default, as a sample size below 20 may not provide enough power to detect significant differences between the sample data and the normal distribution. Outliers were removed by applying the ROUT outlier test. For two-group comparison, datasets were analyzed by unpaired non-parametric two-tailed Mann-Whitney test. For multiple comparisons, we used non-parametric Kruskal-Wallis test followed by uncorrected Dunn’s test. To compare the mean difference between groups with two independent variables two-way ANOVA with Fisher’s least significant difference (LSD) was used. Nested two-sided t-test or nested one-way ANOVA followed by uncorrected Fisher’s LSD test was applied when analyzing technical replicates. Statistical significance was assumed at p < 0.05.

## Results

### HFpEF is associated with macrophage accumulation in RV tissue in mice

To investigate myeloid cells in HFpEF, we employed the established two-hit mouse model for HFpEF (L-NAME, 60% high-fat diet) in *C57BL/6J* mice and dissected cardiac tissue from HFpEF mice in LV (including septum) and RV samples to assess leukocyte counts (**Fig. 1A**)^26^. Total CD45^+^ leukocyte counts trended higher in HFpEF-RVs compared to HFpEF-LVs, mainly due to expansion of cardiac CD64^+^ macrophages (**Fig. 1B-E, extended data figure 1A, B**). To explore a possible association of macrophage expansion with RVD in mice undergoing the two-hit model, we assessed RV function, yet detected no signs of RV hypertrophy with systolic RV pressures in the physiological range (**Extended data fig. 1C**).

**Fig. 1.**
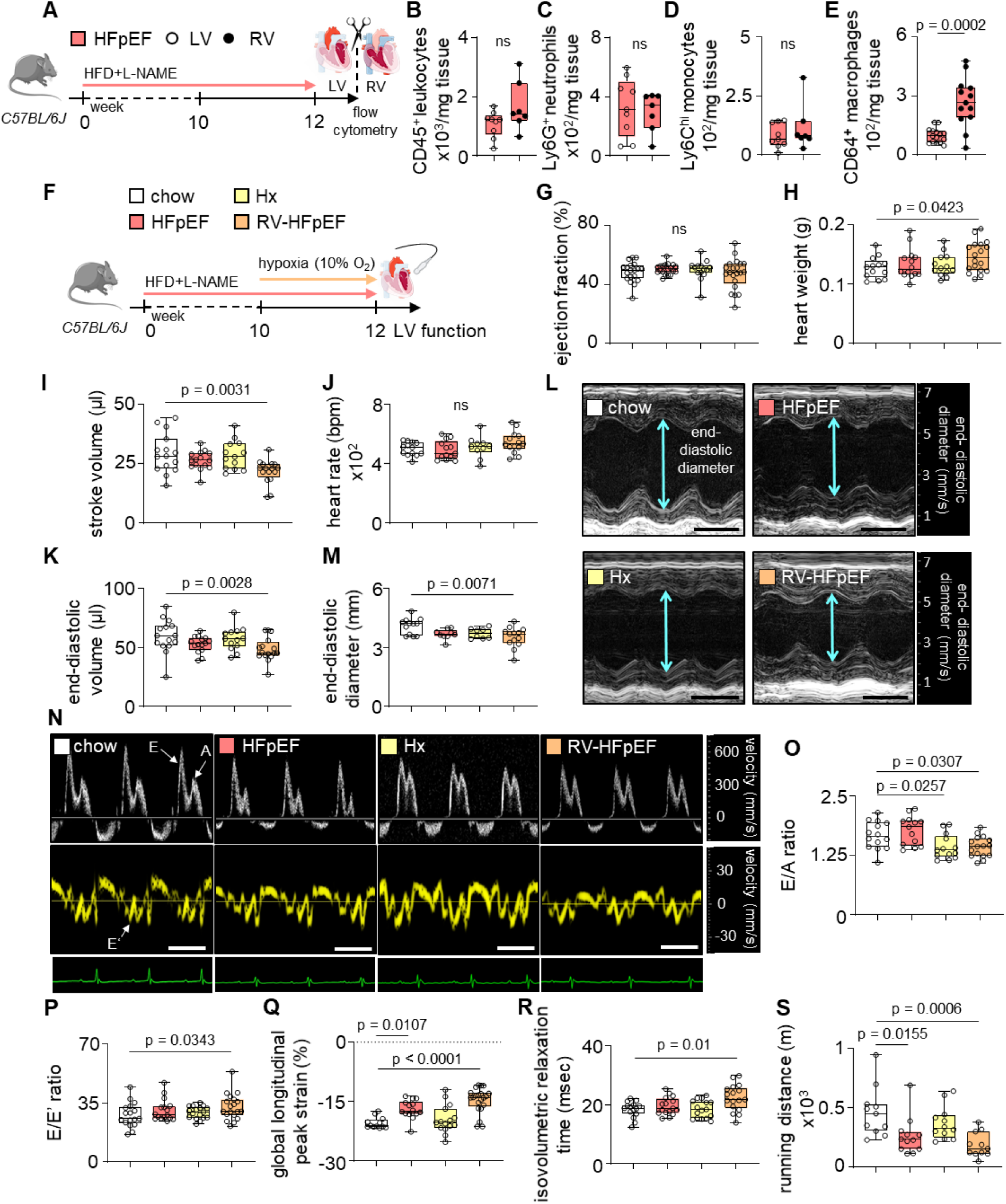
Chronic hypoxia exacerbates LV diastolic dysfunction in HFpEF. **A,** Experimental outline of HFpEF mice induced by treatment of *C57BL/6J* mice with L-NAME and 60% HFD. **B, C, D, E,** Flow cytometric analyses of LV and RV processed tissue samples from HFpEF mice: **B**, CD45^+^ leukocytes. **C**, Ly6G^+^ neutrophils. **D**, Ly6C^hi^ monocytes and **E**, CD64^+^ macrophages (n=7-15). A Mann–Whitney test was used. **F**, Experimental outline of *C57BL/6J* mice treated with i) chow, ii) HFpEF (L-NAME, 60% HFD), iii) chronic hypoxia (Hx, 10% O_2_) or iv) HFpEF and hypoxia (RV-HFpEF). **G**, Echocardiographic assessment of ejection fraction in chow, HFpEF, Hx and RV-HFpEF (n=14-21) mice. **H**, Heart weight of chow, HFpEF, Hx and RV-HFpEF (n=13-18) mice. **I**, Echocardiographic assessment of stroke volume (n=13-16), **J**, Heart rate (n=10-14) and **K**, end-diastolic left ventricular volume in chow, HFpEF, Hx and RV-HFpEF (n=13-16) mice**. L**, Representative echocardiographic M-mode images of the LV in the parasternal long-axis view indicated with blue arrows. scale bar=1 ms. **M**, Analyses of end-diastolic LV diameter of chow, HFpEF, Hx and RV-HFpEF (n=8-13) mice. **N**, Representative echocardiographic pulsed-wave Doppler tracings to calculate the early (E) and late (A) wave peak velocities indicated with white arrows (top panel) and tissue Doppler tracings to calculate early filling velocity (E′) indicated with white arrow (middle panel). Scale bar=1 ms. **O**, E/A ratio (n=13-16), **P**, E/E’ ratio (n=17-19), **Q**, Global longitudinal peak strain (n=12-21) and **R**, isovolumic relaxation time in chow, HFpEF, Hx and RV-HFpEF (n=15-17) mice. **S**, Running distance of chow, HFpEF, Hx and RV-HFpEF (n=11-12) mice in exercise exhaustion test. Kruskal-Wallis test followed by uncorrected Dunn’s multiple comparisons test was used. Data are presented as median with interquartile range (IQR). HFpEF, heart failure with preserved ejection fraction. LV, left ventricle. RV, right ventricle. HFD, high-fat diet. L-NAME, Nω-nitro-L-arginine methylester hydrochloride. ns, not significant.

### Oxygen deprivation fuels LV diastolic dysfunction

A substantial hurdle for studies on RVD is the lack of suitable small animal models^21^, as mice are generally protected from end-stage RV failure and develop only moderate PH and RV hypertrophy^27^. Hence, we designed a novel HFpEF mouse model associated with RVD by complementing the two-hit HFpEF model with an additional risk factor for pulmonary hypertension and RVD development, namely exposure to chronic hypoxia, to induce manifest RVD in mice undergoing the two-hit HFpEF model. Of note, hypoxemia is a common finding in HFpEF patients with secondary PH, in particular during exercise^28–30^. Moreover, chronic hypoxia represents an important accelerator of proinflammatory and profibrotic processes, as Ly6C^lo^ monocytes were shown to sense hypoxia in maladaptive tissue remodeling^31^. Based on this evidence, we hypothesized that reduced oxygen levels in combination with concomitant metabolic and hypertensive stress would fuel RV remodeling and RVD development in the two-hit HFpEF model (henceforth referred to as the ‘RV-HFpEF’ model, **Fig. 1F**).

Exposure to chronic hypoxia (10% O_2_) for two weeks starting at week 10 after HFpEF induction in mice resulted in preserved EF, reduced stroke volumes, and concentric LV hypertrophy, evident as increased heart weights, reduced LV end-diastolic diameters and volumes (**Fig. 1G-M**). RV-HFpEF mice presented with pronounced LV diastolic dysfunction indicated by decreased E/A ratios, increased E/E’ ratios, reduced global longitudinal peak strain, and increased isovolumetric relaxation time when compared to controls (**Fig. 1N-R**). Of note, exposure to chronic hypoxia alone induced signs of diastolic dysfunction independent of underlying HFpEF, confirming that oxygen depletion contributes to LV diastolic dysfunction independent of metabolic stress (**Fig. 1N-R**). Plasma NT-proBNP levels, RV gene expression of myosin heavy chain 7 (Myh7) and the natriuretic peptide B (Nppb), and hepatic triglycerides were increased in RV-HFpEF mice (**extended data fig. 1D-F**), suggestive of an aggravated HFpEF phenotype. Exercise intolerance is a cardinal symptom of HFpEF patients, which we diagnosed in our animal model by shorter running distances of HFpEF mice when compared to normoxic controls in an established treadmill test (**Fig 1S**). Impaired LV function in RV-HFpEF mice relative to HFpEF mice is partially counteracted by increased hemoglobin content and hematocrit (**Extended data fig. 1G, H**), resulting in an overall similar exercise performance. In sum, reduced oxygen availability exacerbates LV diastolic dysfunction.

### Chronic hypoxia induces RV remodeling and dysfunction in HFpEF mice

RV phenotyping revealed RV hypertrophy in hypoxia-treated groups, independent of underlying HFpEF, evident as increased RV weights and Fulton’s index (**Fig. 2A-D**). At the functional level, we found decreased pulmonary artery peak velocities, elevated RV systolic pressures, and reduced echocardiographically-assessed tricuspid annular plane systolic excursions in RV-HFpEF mice, altogether indicating manifest RVD in RV-HFpEF mice (**Fig. 2E-J**).

**Fig. 2.**
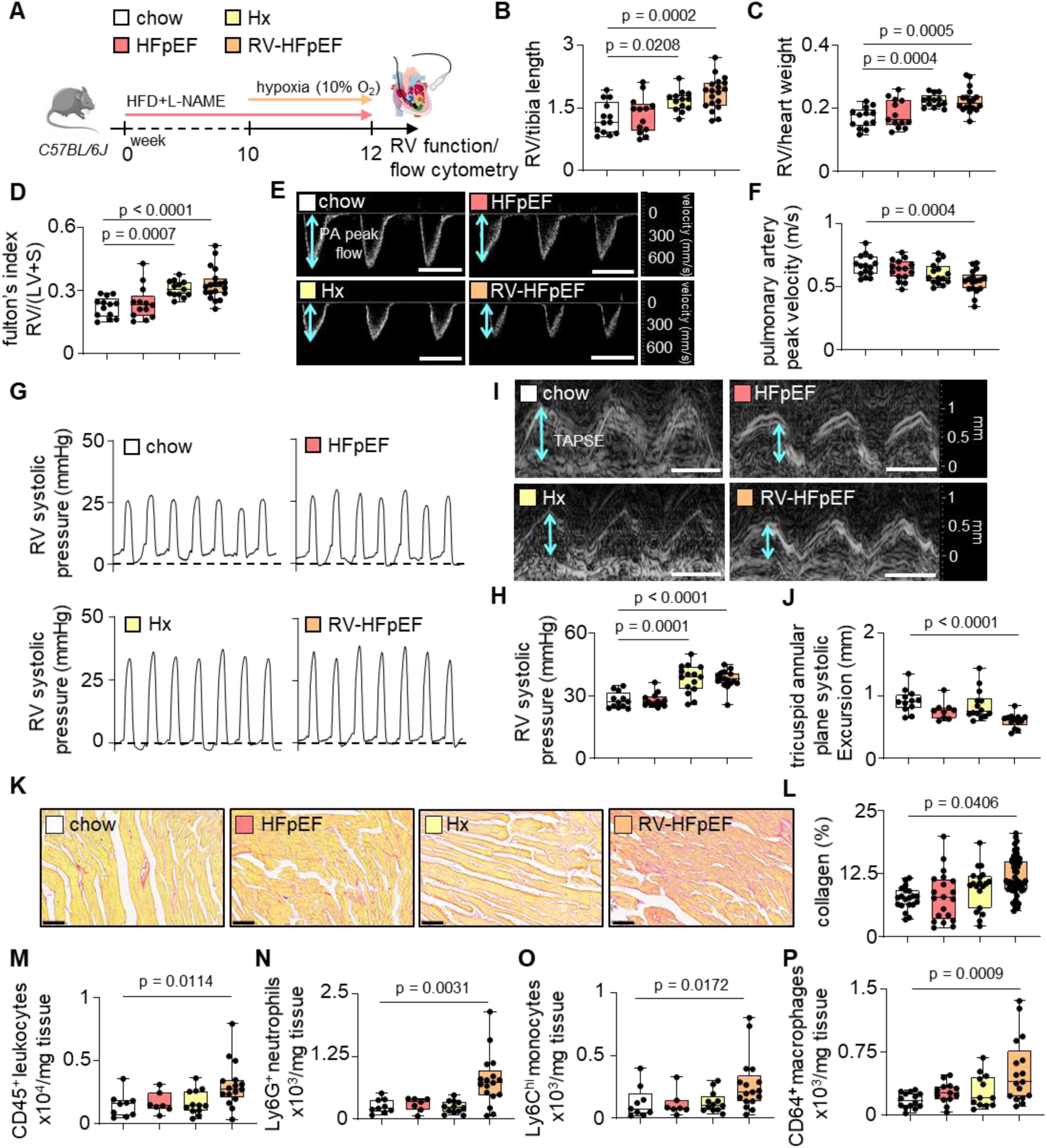
Combined hypoxia and HFpEF results in RVD. **A,** Experimental outline of *C57BL/6J* mice treated with i) chow, ii) HFpEF (L-NAME, 60% HFD), iii) chronic hypoxia (Hx, 10% O_2_) or iv) HFpEF and hypoxia (RV-HFpEF). **B,** RV weight-to-tibia length ratio (RV/tibia length, n=13-19), **C**, RV weight-to-heart weight ratio (RV/heart weight, n=12-19) and **D,** fulton’s index (RV mass relative to LV and septal mass ((RV/(LV+S)) in chow in chow, HFpEF, Hx and RV-HFpEF (n=13-19) mice. **E**, Representative echocardiographic pulsed-wave Doppler tracings to calculate pulmonary artery peak velocity indicated with blue arrows. scale bar=1 ms. **F**, Quantitative analyses of pulmonary artery peak velocity profiles in chow, HFpEF, Hx and RV-HFpEF (n=15-17) mice. **G**, Representative RV systolic pressure profiles. **H**, Quantitative analyses of RV systolic pressures in chow, HFpEF, Hx and RV-HFpEF (n=12-16) mice. **I**, Representative echocardiographic M-mode images to calculate tricuspid annular plane systolic excursion (TAPSE) indicated with blue arrows. scale bar=1 ms. **J**, Quantitative analyses of TAPSE in chow, HFpEF, Hx and RV-HFpEF (n=9-14) mice. **K,** Representative images of picrosirius red staining of RV tissue sections. scale bar=50 µm. **L**, Quantitative analyses of collagen content in RV sections of chow, HFpEF, Hx and RV-HFpEF (n=20-55). Each dot represents a field of view. A nested one-way ANOVA followed by uncorrected Fisher’s least significant difference test was used. **M** Flow cytometric quantification of RV CD45^+^ leukocytes, **N**, Ly6G^+^ neutrophils, **O**, Ly6C^hi^ monocytes and **P**, CD64^+^ macrophages of chow, HFpEF, Hx and RV-HFpEF (n=7-17) mice. Kruskal-Wallis test followed by uncorrected Dunn’s multiple comparisons test was used. Data are presented as median with interquartile range (IQR). HFpEF, heart failure with preserved ejection fraction. LV, left ventricle. RV, right ventricle. HFD, high-fat diet. L-NAME, Nω-nitro-L-arginine methylester hydrochloride.

Women represent a greater proportion of HFpEF patients when compared to HF with reduced EF (HFrEF) and outnumber men by a ratio of 2:1^32,33^. In contrast, female mice exposed to the two-hit HFpEF model were protected from LV diastolic dysfunction^34^. Differently from findings in the two-hit model, we did not observe sex differences in LV or RV outcomes in our novel RV-HFpEF mouse model (**Extended data fig. 2A-J**), matching the clinical scenario where female sex is one of the strongest risk factors for pulmonary arterial hypertension (PAH) and associated RVD development^35^.

Next, we detected increased gene expression of pro-fibrotic markers and substantial collagen deposition in RV tissues from RV-HFpEF mice, but not in groups exposed to HFpEF or chronic hypoxia alone, indicating that a combination of concomitant metabolic, hypertensive and hypoxic stress is necessary to induce substantial RV remodeling (**Fig. 2K, L, extended data fig. 2K**). To test, whether myeloid cells contribute to RV remodeling in RV-HFpEF, we performed multi-color flow cytometry of RV tissues and found elevated CD45^+^ leukocyte, Ly6G^+^ neutrophil, Ly6C^hi^ monocyte, and resident CD64^+^ macrophage counts relative to controls (**Fig. 2M-P**). Of note, immune cell accumulation in RV tissue was not triggered by chronic hypoxia alone, but only became evident when combining metabolic stress, hypertension <u>and</u> chronic hypoxia (**Extended data fig. 3A-E**), underscoring the relevance of concomitant systemic stressors for RVD development in HFpEF. In addition, RV-HFpEF reduced the percentage of resident macrophage subsets (**Extended data fig. 3F, G**) and increased cardiac T-cell, B-cell and natural killer cell counts (**Extended data fig. 3H-K**), indicating fundamental dysregulation of leukocyte dynamics in RVD associated to HFpEF.

### Dysregulated innate immune responses and extracellular matrix disorganization denote key pathophysiological features of RV tissue remodeling in HFpEF

To understand the molecular mechanisms involved in RV remodeling in HFpEF, we performed unbiased mass spectrometry-based proteomic profiling of bulk RV tissue from all experimental groups. Single-sample gene set enrichment analysis (ssGSEA) of Reactome pathways revealed upregulation of innate immune system and extracellular matrix organization processes in RV tissue from RV-HFpEF mice relative to other groups (**Fig. 3A, B**), confirming our initial findings from flow cytometry and histology. Importantly, these pathways remained significantly enriched when comparing RV-HFpEF proteomes with those from HFpEF-only or hypoxia-only mice, indicating that innate immune cell activation and extracellular matrix deposition are a consequence of the combined systemic stressors in RV-HFpEF. Enriched proteins from these pathways can be classified by function as follows: i) cell adhesion molecules such as integrin beta 2 (ITGB2), integrin alpha M (ITGAM), and cluster of differentiation 44 (CD44), ii) strong inflammatory mediators such as the alarmins S100 calcium-binding protein A8 (S100A8) and S100 calcium-binding protein A9 (S100A9), and iii) matricellular proteins such as versican (VCAN), secreted protein, acidic, rich in cysteine (SPARC) and fibulin-2 (FBLN2) (**Fig. 3C-E**). In addition, proteomic profiling also revealed increased abundance of proteins associated with platelet activation and the adaptive immune system in RV-HFpEF mice (**Fig. 3 A, B**), both pathophysiological features in clinical HFpEF^36–40^. Furthermore, RV tissues from RV-HFpEF mice presented with proteomic signatures indicative of blunted mitophagy, reduced glycolysis, and impaired mitochondrial metabolism, all of which represent pathophysiological hallmarks previously described in HFpEF myocardial tissue^41,42^.

**Fig. 3.**
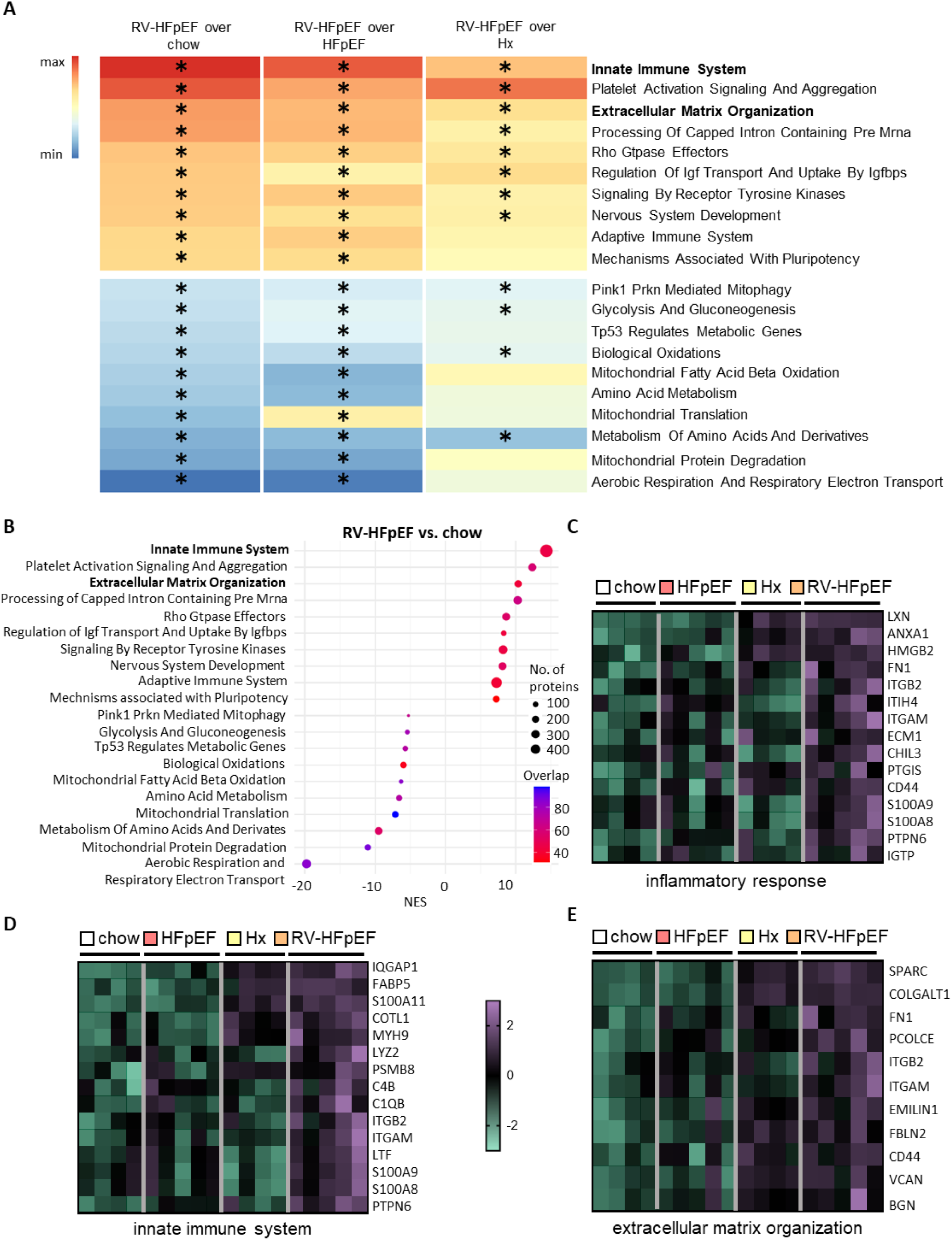
Extracellular matrix turnover and innate immune system activation in RVD associated with experimental HFpEF. **A**, Heatmap showing the top and bottom 10 Reactome pathways identified by single-sample Gene Set Enrichment Analysis (ssGSEA) for RV-HFpEF versus chow, together with the enrichment scores of these pathways in the other experimental comparisons (HFpEF and hypoxia (Hx)). Pathway enrichment was calculated from log₂ fold-change values (red, upregulated; blue, downregulated). Asterisks indicate significantly enriched pathways (FDR < 0.05) **B**, Fold enrichment dot plot showing the top and bottom 10 Reactome pathways identified by single-sample Gene Set Enrichment Analysis (ssGSEA) for RV-HFpEF versus chow. A positive Normalized Enrichment Score (NES) indicates enrichment of the respective pathway in RV-HFpEF relative to chow with a negative NES indicating the same for chow vs. RV-HFpEF. The size of each dot represents the total number of proteins annotated to each biological process in the Reactome database and the colors of the plots indicate the number of proteins from the set contributing to a specific enrichment for a given pathway. NES were calculated with Kolmogorov-Smirnov-like running sum statistics. **C**, **D**, **E** Heatmap of expression values of the top upregulated proteins in RV tissue of RV-HFpEF vs. chow (logFC>1; p<0.05; z-scored) belonging to the respective Reactome pathways (inflammatory response, innate immune system, extracellular matrix organization). HFpEF, heart failure with preserved ejection fraction. RV, right ventricle. Hx, hypoxia. * p<0.05.

### HFpEF-RVs and HFpEF-LVs exhibit distinct ventricular remodeling responses when exposed to systemic stressors

Cumulative evidence suggests that LV and RV fundamentally differ regarding their molecular, cellular and physiological adaptive response to adverse loading and systemic stressors^43^. To understand the role of myeloid cells in biventricular dysfunction due to systemic metabolic, hypertensive and hypoxic stress, we directly compared LV and RV tissue remodeling responses employing the RV-HFpEF model (**Fig. 4**). Interestingly, we found more pronounced collagen deposition in RVs accompanied by marked accumulation of total CD45^+^ leukocytes, Ly6C^hi^ monocytes and CD64^+^ macrophages when compared to LVs from the same animals (**Fig. 4A-F**). A similar pattern was evident for monocyte-derived CCR2^+^ macrophage subsets, which we found in higher numbers in RVs when compared to LVs, suggesting that the proportion of macrophages originally recruited from the blood pool of circulating monocytes is higher in RVs compared to LVs from RV-HFpEF animals, as the chemokine receptor CCR2 is preferentially expressed by recruited myeloid subsets^44^ (**Fig. 4G, H**). The notion of ventricle-specific pathophysiological responses to systemic stressors was further supported by direct comparison of regulated pathways at the proteomic level. In RV-HFpEF, proteins associated with innate immune system activation, extracellular matrix dysregulation, and cellular adhesion were significantly enriched in RV samples when compared to LV samples (**Extended data fig. 4A, B).** Moreover, RVs were enriched in proteins involved in leukocyte and macrophage chemotaxis compared to LVs (**Fig 4I, extended data fig. 4C**), prompting experiments to decipher the origin and fate of RV macrophages.

**Fig. 4.**
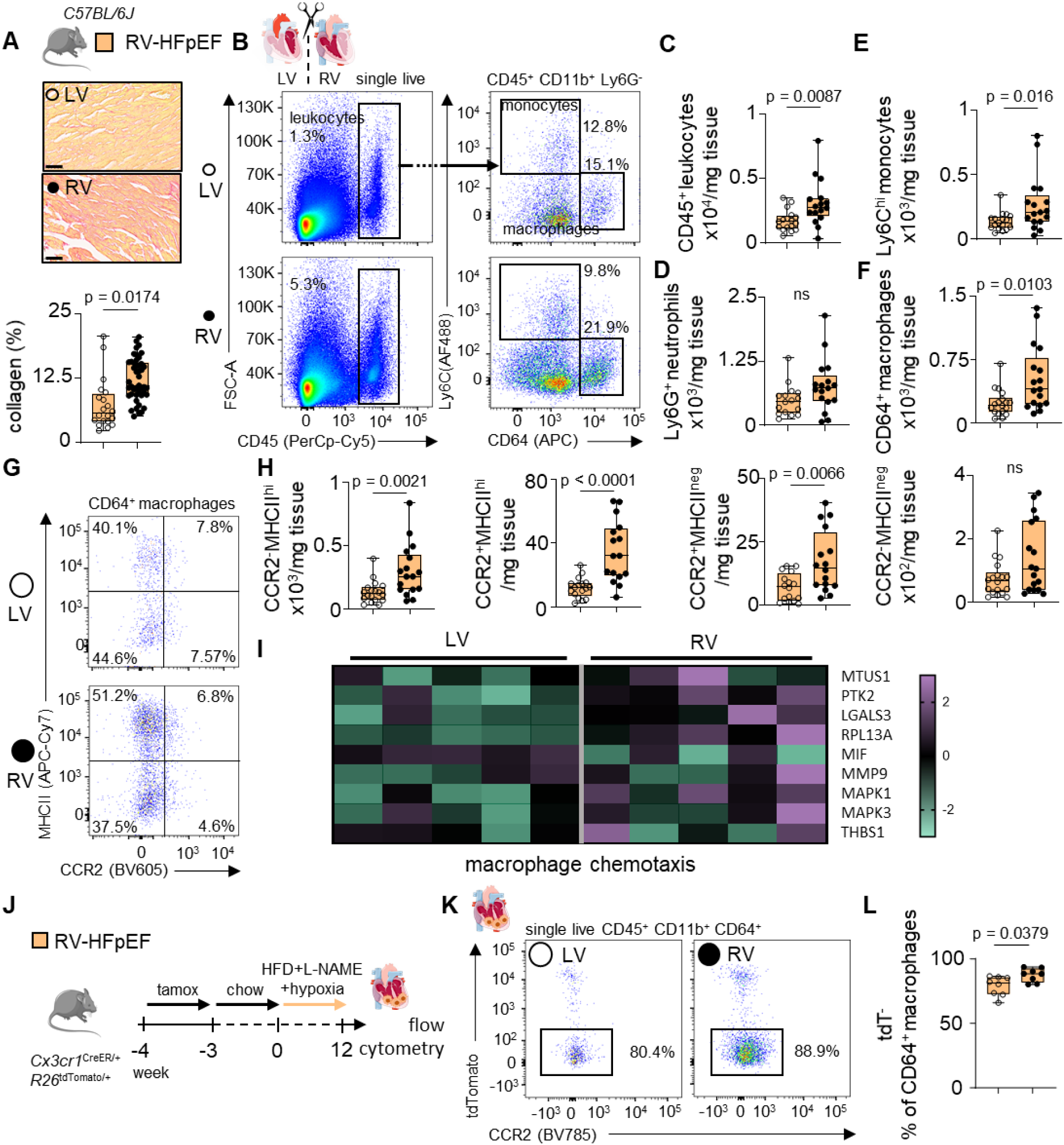
RV inflammation, myeloid cell recruitment and remodeling in RV-HFpEF mice. **A**, Representative images of picrosirius red staining. scale bar=50 µm. Quantitative analyses of collagen content in LV and RV (n=20-45) tissues from RV-HFpEF mice. Each dot represents a field of view. A nested t test was used. **B**, Representative flow plots of LV and RV leukocytes and myeloid cells in HFpEF mice exposed to hypoxia (10% O_2_, RV-HFpEF). **C**, Comparison of LV and RV CD45^+^ leukocytes, **D**, Ly6G^+^ neutrophils, **E**, Ly6C^hi^ monocytes and **F**, CD64^+^ macrophages in RV-HFpEF mice (n=16-18). **G**, Representative flow plots after gating for LV and RV macrophage subsets and **H**, corresponding quantification of CCR2^-^ MHCII^neg^, CCR2^-^ MHCII^hi^, CCR2^+^ MHCII^neg^ and CCR2^+^ MHCII^hi^ subset in LV and RV (n=16-18) of RV-HFpEF mice. **I**, Heatmap of z-scored protein expression values in RV versus LV tissue from RV-HFpEF mice for proteins annotated to the Gene Ontology term “macrophage chemotaxis. **J**, Experimental outline of Fate-mapping in Cx3cr1^CreER/+^R26^tdTomato/+^ RV-HFpEF mice treated with tamoxifen-rich diet (tamox) for 1 week prior to a 3-week flushout followed by HFpEF (L-NAME, 60% HFD) and chronic hypoxia (Hx, 10% O_2_). **K**, Representative flow plots of LV and RV tdTomato negative (tdT^-^) macrophages in RV-HFpEF. **L**, Comparison of LV and RV (n=8) recruited tdT^-^ macrophages as a percentage of CD64^+^ macrophages. Mann–Whitney test was used. Data are presented as median with interquartile range (IQR). HFpEF, heart failure with preserved ejection fraction. LV, left ventricle. RV, right ventricle. HFD, high-fat diet. L-NAME, Nω-nitro-L-arginine methylester hydrochloride. Tamox, tamoxifen-rich diet. ns, not significant.

### Monocyte recruitment is the major source of RV myeloid cell expansion in HFpEF-associated RVD

Understanding origin and fate of immune cells has direct translational implications, as cells being causally involved in cardiac pathogenesis have been previously thought to be diagnostically inaccessible within the heart, but may be in reality quantifiable – and therefore targetable – in easily accessible peripheral blood samples. To explore the origin of macrophage expansion in RV-HFpEF ventricles, we first performed Ki67 staining on CD64^+^ macrophages in both ventricles from RV-HFpEF mice (**Extended data fig. 5A-B**) to probe whether CD64^+^ macrophage expansion was attributable to proliferation of cardiac macrophages. In both ventricles, CD64^+^ macrophages presented with strongly elevated proliferation rates in response to RV-HFpEF, with no differences between LVs and RVs. As proliferation rates appeared comparable between the ventricles, we hypothesized that myeloid cell recruitment differed between them and represented the primary source of macrophage accumulation in RV tissue. To test this, we used *Cx3cr1^CreER/+^R26^tdTomato/+^* mice for genetic fate mapping experiments in our novel RV-HFpEF model. In this reporter strain, *Cx3cr1*-expressing myeloid cells are efficiently labeled with tdTomato upon tamoxifen exposure (**Extended data fig. 5C-E**). In RV-HFpEF mice, myeloid cell recruitment expanded so that over 80 % of ventricular macrophages derived from circulating monocytes, an increase compared to the steady state (data not shown). Of note, myeloid cell recruitment was significantly increased in RV tissue compared to LV tissue from the same RV-HFpEF animals (**Fig. 4J-L**). Viewed together, these data indicate that myeloid cell recruitment becomes the primary source of cardiac macrophages in HFpEF, which is exacerbated in RV tissue when compared to LV tissue.

### Myeloid cells propel RVD in HFpEF via sterile inflammation

To decipher how macrophage accumulation contributes to RV remodeling in HFpEF, we next performed gene expression profiling in bulk RV tissue, MACS-sorted CD64^+^ macrophages and Ly6G^+^ neutrophils from RV tissue of RV-HFpEF mice, screening for pro-inflammatory and pro-fibrotic transcripts (**Fig. 5A**). We found inflammatory mediators such as tumor necrosis factor alpha and interleukin 1 beta particularly enriched in neutrophils from RV-HFpEF mice, while pro-fibrotic marker expression was associated with CD64^+^ macrophages, highlighting again the importance of innate immune cell populations in RV tissue remodeling in HFpEF.

**Fig. 5.**
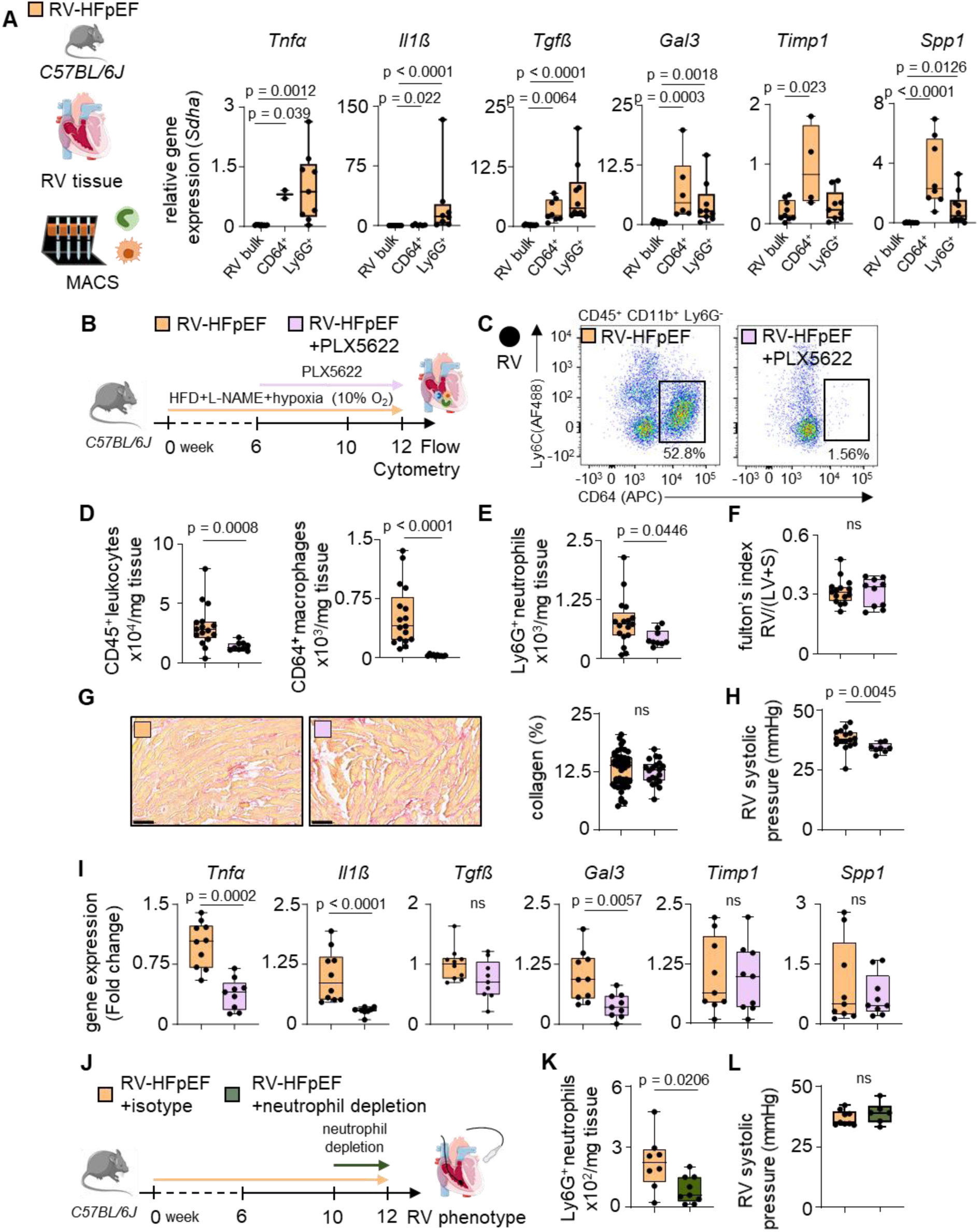
Macrophage depletion improves RV function in RV-HFpEF mice. **A**, Quantitative PCR of RV-HFpEF mice. *Tnf*α, *Il1ß*, *Tgfß*, *Gal3*, *Timp1* and *Spp1* expression in RV bulk tissue, MACS sorted CD64^+^ macrophages and Ly6G^+^ neutrophils (n=2-10). Gene expression was normalized to *Sdha*. Kruskal-Wallis test followed by uncorrected Dunn’s multiple comparisons test was used. **B,** Experimental outline for RV-HFpEF mice treated with colony-stimulating factor 1 receptor (CSF1R) inhibitor (PLX5622) for the last 6 weeks of the experiment (RV-HFpEF+PLX5622). **C**, Representative flow plots showing macrophages from RV-HFpEF mice and RV-HFpEF+PLX5622 mice. **D**, Corresponding quantification of RV CD45^+^ leukocytes in RV-HFpEF and RV-HFpEF+PLX5622 (n=9-16) mice (left) and CD64^+^ macrophages in RV-HFpEF and RV-HFpEF+PLX5622 (n=10-17) mice (right). **E**, Quantification of Ly6G^+^ neutrophils in RV-HFpEF and RV-HFpEF+PLX5622 (n=10-17) mice. **F,** fulton‘s index (RV weight to LV + septum weight (RV/(LV+S)) from RV-HFpEF and RV-HFpEF+PLX5622 (n=10-15) mice. **G**, Representative images of Picrosirius red staining (left). scale bar=50 µm. Quantitative analyses of RV-HFpEF RVs and RV-HFpEF+PLX5622 (n=20-45) collagen content (right). Each dot represents a field of view. Nested t test was used. **H**, RV systolic pressure of RV-HFpEF and RV-HFpEF+PLX5622 (n=8-16) mice. **I**, Quantitative PCR of RV-HFpEF and RV-HFpEF+PLX5622 mice. *Tnfα* (n=9-10), *Il1ß* (n=8-10), *Tgfß* (n=9-10), *Gal3* (n=9-10), *Timp1* (n=9) and *Spp1* expression in RV bulk tissue of RV-HFpEF and RV-HFpEF+PLX (n=9-10). Expression values were calculated relative to *Sdha* expression using the 2^−ΔΔCt^ method. **J**, Experimental outline for RV-HFpEF mice treated with neutrophil depletion (RV-HFpEF+neutrophil depletion) or control (RV-HFpEF+isotype) for the last 2 weeks of the experiment. **K**, Quantification of Ly6G^+^ neutrophils in RV-HFpEF+isotype and RV-HFpEF+neutrophil depletion (n=8-9). **L**, RV systolic pressure of RV-HFpEF+isotype and RV-HFpEF+neutrophil depletion mice (n=6-9). Mann–Whitney test was used. Data are presented as median with interquartile range (IQR). HFpEF, heart failure with preserved ejection fraction. RV, right ventricle. HFD, high-fat diet. L-NAME, Nω-nitro-L-arginine methylester hydrochloride. ns, not significant.

Based on these data, we hypothesized that macrophages with a pro-fibrotic profile may be particularly relevant to RV remodeling and subsequent RVD development in HFpEF. To test this notion, we depleted the cardiac macrophage pool by pharmacologically targeting the colony-stimulating factor 1 receptor (CSF1R)^45^. CSF1R promotes macrophage differentiation, survival, and myeloid cell proliferation^46^, whereas CSF1R inhibition rapidly results in macrophage depletion of cell populations relying on CSF1R^45^. Macrophage depletion in RV-HFpEF mice was achieved by treatment with the CSF1R inhibitor PLX5622 for six weeks, starting six weeks after HFpEF induction (**Fig. 5B**). This experimental design was chosen based on the hypothesis that potential pro-fibrotic effects mediated by myeloid cells may only become effective at a later disease stage at the time of RVD development and when LV diastolic dysfunction is already established. Treatment with PLX5622 sufficiently depleted cardiac CD64^+^ macrophages below 2% of total cardiac leukocytes whereas blood leukocyte counts and LV function remained stable (**Fig. 5C, D, extended data fig. 6A-G, extended data fig. 7A-C**). Of note, we also found cardiac neutrophil counts reduced in RV tissue from macrophage-depleted RV-HFpEF mice, suggesting macrophage-neutrophil interaction in HFpEF-associated RVD (**Fig. 5E**). While RV remodeling, as evidenced by RV hypertrophy and RV matrix deposition remained unchanged in response to macrophage depletion, RV hemodynamics, pro-inflammatory and pro-fibrotic gene expression patterns were rescued in RV-HFpEF mice upon macrophage depletion (**Fig. 5F-I**). To probe for potential indirect effects of macrophages on RV tissue via macrophage-neutrophil interactions, we employed a neutrophil depletion strategy using antibodies directed against neutrophil surface markers (**Fig. 5J**). Neutrophil depletion effectively reduced neutrophil abundance in RV tissue in RV-HFpEF mice (**Fig. 5K**), yet failed to rescue RV hemodynamics, RV function and RV inflammatory profiles (**Fig. 5L, extended data fig. 8A-D).**

Our results indicate that macrophage depletion partially rescues RV phenotypes in RV-HFpEF mice, suggesting that myeloid cells are causally involved in RVD pathogenesis in HFpEF by directly mediating tissue inflammation and remodeling. Conversely, our results also reveal RV remodeling processes that are independent of myeloid cell accumulation in HFpEF associated RVD.

### RV dilation is associated with unique blood leukocyte signatures in HFpEF patients

To assess the relevance of our findings in the clinical setting, we retrospectively studied leukocyte counts in a consecutive HFpEF patient cohort recruited by our Collaborative Research Center 1470 (n=123, further referred to as ‘CRC1470 cohort’), which were included according to the ACC guidelines^47^ matching the following criteria: i) EF>50%, ii) N-terminal pro-B-type natriuretic peptide (NT-proBNP) >125 pg/mL, iii) New York Heart Association (NYHA) stage II or III, and iv) biventricular echocardiography (**Fig. 6A**). We dichotomized the CRC1470 patient cohort by RV dilation in HFpEF patients with RV dilation (>41mm) and HFpEF patients without RV dilation (<41mm). RV dilation is often associated with RVD and worse outcomes and according to a meta-analysis is the most sensitive measure predictive of mortality in HFpEF patients^48–50^. According to this stratification, 23% of HFpEF patients had RV dilation (**Fig. 6B, C**). The patient characteristics revealed that HFpEF patients with RV dilation were more frequently male, had higher NT-proBNP serum levels and had experienced atrial fibrillation more frequently compared to HFpEF patients with regular RV diameters (**Table 1**). Average sPAP, sPAP-TAPSE ratio, and right atrial dilation were all impaired in HFpEF patients with RV dilation, indicating signs of RVD. Interestingly, total leukocyte counts were significantly reduced in HFpEF patients with RV dilation when compared to HFpEF patients with normal RV function (**Table 1**). White blood cell differentials revealed a myeloid cell bias towards increased percentages of monocytes in HFpEF patients with RV dilation (**Fig. 6D**, **Table 1**), whereas neutrophils and lymphocyte counts remained comparable in both groups (**Table 1**).

**Fig. 6.**
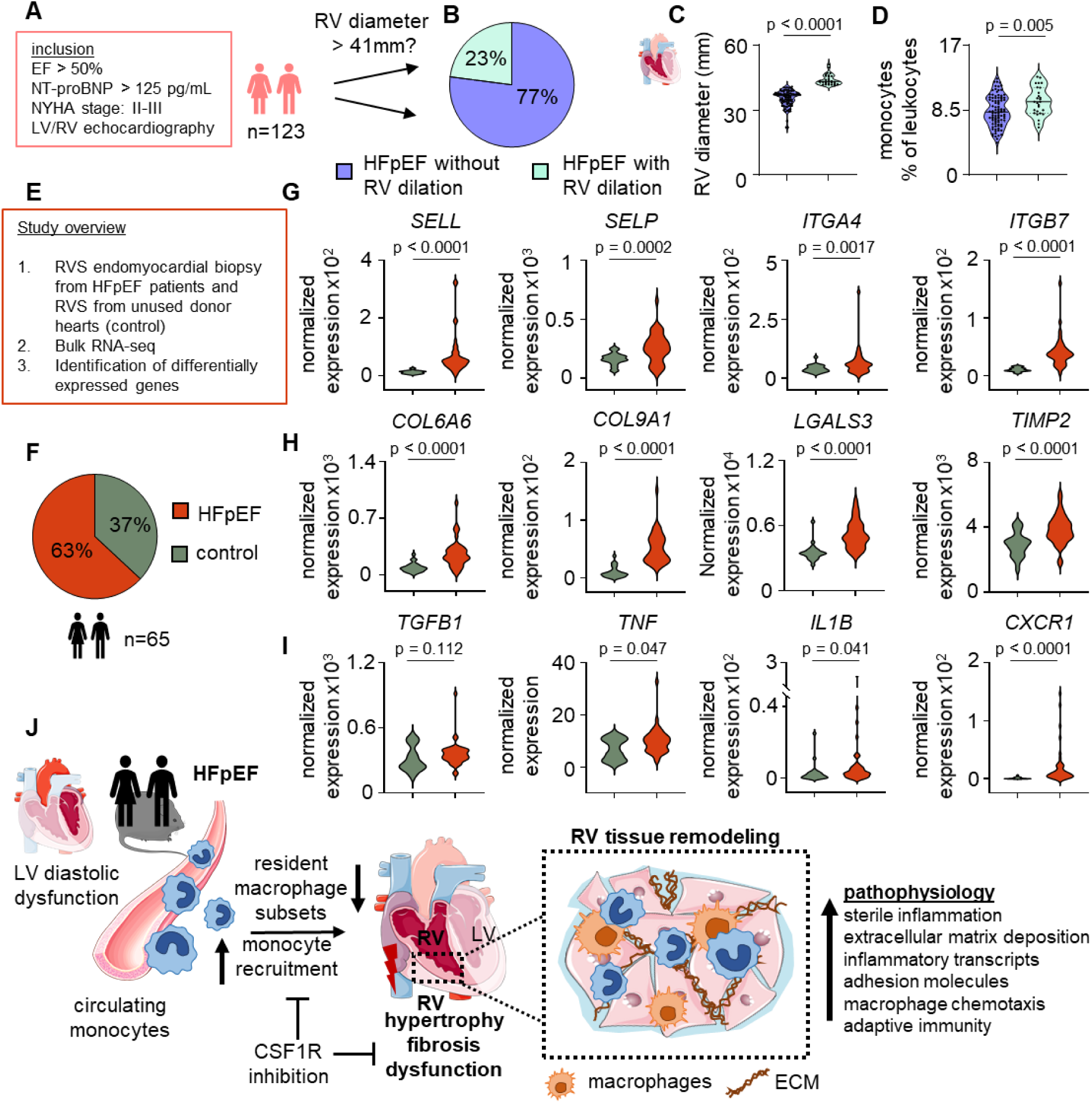
HFpEF patients show increased circulating macrophages and elevated extracellular matrix turnover and inflammation in RV biopsies. **A,** Experimental outline of patient cohort with HFpEF from Collaborative Research Center (CRC) 1470 (n=123). **B**, Percentage of HFpEF patients with and without RV dilation. **C**, Echocardiographic analysis of RV diameter. **D**, Percentage monocytes of leukocytes in HFpEF with RV dilation and HFpEF without RV dilation (n=28-95). **E**, Study overview for the identification of differentially expressed genes in RV septum (RVS) biopsies from HFpEF vs. controls using DESeq2. **F**, Percentage of HFpEF and control patients included for bulk RNA-seq of RVS biopsies. **G**, **H**, **I**, Differential expression of selected adhesion molecules (**G**), pro-fibrotic markers (**H**) and inflammatory markers (**I**) in RVS biopsies from control (n=24) vs. HFpEF (n=41) patients. P values were adjusted for multiple comparisons by using the Benjamini-Hochberg method. **J**, Graphical abstract. HFpEF, heart failure with preserved ejection fraction. CSF1R, Colony stimulating factor 1 receptor.

**Table 1.** Baseline characteristics of CRC1470 patient cohort.

| Characteristics | HFpEF<br>Without RV dilation | HFpEF with RV<br>dilation | P-value |
| --- | --- | --- | --- |
| **HFpEF** |  |  |  |
| N (%) | 95 (77%) | 28 (23%) |  |
| Age (years) | 75 (50, 87) | 76 (61, 89) | 0.886 |
| Female gender (%) | 55 (58%) | 9 (32%) | 0.017 |
| Male gender (%) | 40 (42%) | 19 (68%) |  |
| NYHA class II (%) | 66 (69%) | 18 (64%) | 0.6 |
| NYHA class III (%) | 29 (31%) | 10 (36%) |  |
| NT-pro-BNP (pg/mL) | 395 (128, 2,412) | 499 (154, 2,047) | 0.062 |
| LV EF (%) | 57.0 (45.0, 74.0) | 54.0 (50.0, 66.0) | 0.11 |
| **RV function** |  |  |  |
| RV diameter | 36.64 (22.00, 40.90) | 43.25 (41.41, 51.23) | <0.0001 |
| sPAP (mmHg) | 31 (16, 72) | 38 (20, 57) | 0.002 |
| TAPSE-sPAP Ratio | 0.74 (0.37, 1.69) | 0.59 (0.29, 1.20) | 0.012 |
| RA dilation | 47 (53%) | 21 (88%) | 0.002 |
| **Comorbidities** |  |  |  |
| Hypertension | 75 (79%) | 23 (82%) | 0.7 |
| Atrial Fibrillation | 54 (57%) | 24 (86%) | 0.005 |
| Diabetes Mellitus<br>Type II | 20 (21%) | 5 (18%) | 0.7 |
| Congestion (Edema) | 23 (25%) | 9 (32%) | 0.4 |
| **Inflammation** |  |  |  |
| Leukocyte counts<br>(x10<sup>3</sup>/μl) | 7.00 (3.82, 13.80) | 6.56 (3.19, 9.62) | 0.029 |
| Monocyte % | 8.20 (4.20, 12.60) | 9.60 (6.20, 12.90) | 0.005 |
| Neutrophil % | 65 (44, 83) | 65 (45, 78) | 0.5 |
| Lymphocyte % | 24 (5, 41) | 22 (13, 41) | 0.2 |

HFpEF, Heart failure with preserved ejection fraction, NYHA, New York Heart association, NT-pro-BNP, N-Terminal Pro-B-Type Natriuretic Peptide, LVEF, Left ventricular ejection fraction, RV, right ventricle S', TAPSE, tricuspid annular plane systolic excursion, sPAP, systolic pulmonary artery pressure, RA, right atrium CRP. P-values calculated using a Wilcoxon rank sum test, Fisher's exact test or Pearson's Chi-squared test. Data are presented as median with range from min to max or n (%).

Finally, we asked whether cardiac macrophage-associated mechanisms identified in RV-HFpEF mice translate to the clinical scenario. Reanalysis of a published bulk RNA sequencing dataset from RV septum endomyocardial biopsies from unused donor hearts (controls) and HFpEF patients^23^ (**Fig. 6E, F**) revealed i) increased expression levels of cell adhesion molecules associated with leukocyte-endothelial cell interaction such as selectins and integrins (*SELL, SELP, ITGA4, ITGB7*, **Fig. 6G**), ii) elevated expression of matricellular proteins and pro-fibrotic mediators such as collagens, galectins and tissue inhibitors of metalloproteinases (*COL6A6, COL9A1, LGALS3, TIMP2*, **Fig. 6H**), and iii) higher expression of pro-inflammatory cytokines and their receptors (*TNF, IL1B, CXCR1)*, while *TGFB1* was not significantly regulated (**Fig. 6I**), altogether replicating and thus, validating our murine proteomics data from RV-HFpEF mice in RV tissue from HFpEF patients.

## Discussion

Collectively, our data indicate that myeloid cells contribute to the pathogenesis of RVD in HFpEF (**Fig. 6J**). This is an important novel finding as ventricle-specific pathomechanisms have, as of yet, not been addressed in the context of HFpEF, despite the emerging recognition of monocyte-derived macrophages as a pathogenic factor of ventricular dysfunction in HFpEF^7,51^. Here, we identified macrophage chemotaxis, unique cell adhesion molecule profiles, monocyte recruitment and extracellular matrix disorganization as mechanisms at play in RVD associated to HFpEF.

Hypoxemia is a common finding in HFpEF patients, in particular during exercise^28–30^. Although the 10% chronic hypoxia used in our RV-HFpEF model exceeds typical clinical levels, it offers a rigorous proof-of-principle test for synergistic effects of hypoxia and metabolic/hypertensive stress on the RV and still reflects a realistic scenario given the high coexistence of HFpEF and COPD^52,53^. Because COPD is a frequent, prognostically adverse HFpEF comorbidity and nearly half of patients with PH due to left heart disease exhibit overlapping hypoxia-related PH^54^, hypoxemia hypoxia represents a substantial and increasingly recognized clinical burden in HFpEF patients.

Our data from HFpEF patients identified lower leukocyte counts in HFpEF with RV dilation compared to HFpEF with normal RV diameters. Whilst leukocyte counts in HFpEF patients with RVD were lower, white blood cell counts were still in the upper physiological range. Reduced leukocyte counts have been reported before in patients with decompensated heart failure and in patients with idiopathic PAH^55,56^. This observation may be, at least in part, explained by our finding of increased expression of leukocyte adhesion molecules in RV tissue, which is associated with enhanced immune cell recruitment to the site of injury and could therefore explain lower levels of circulating white blood cells. Increased expression of adhesion molecules has been previously reported in clinical and experimental HFpEF^39,57,58^ as well as in PH and RV failure^59,60^, suggesting cell adhesion as a key feature in RVD-HFpEF. Another likely explanation for this finding may be a disturbed production or processing of mature leukocytes in the bone marrow of HFpEF individuals. This thought is supported by recent cumulative evidence, linking the risk of incident HFpEF to Clonal hematopoiesis of indeterminate potential (CHIP)^61–63^. Data to test if HFpEF patients with RVD are more frequently affected by CHIP or other hematopoietic disorders such as myelodysplastic syndrome when compared to HFpEF patients without RVD, are to our knowledge not yet available.

Besides the association of circulating leukocyte counts with RV dilation in HFpEF patients, studying leukocyte dynamics in the clinical scenario is restricted by the limited availability of RV tissue samples from HFpEF patients. CD68⁺ macrophages have been documented in endomyocardial RV septal biopsies from HFpEF patients, with increased macrophage burden correlating with distinct clinical phenotypes and suggesting discrete HFpEF endotypes^23^. This concept is reinforced by single-cell RNA sequencing, which identifies a macrophage subset in HFpEF-RVs characterized by a disease-specific transcriptional signature relative to controls^64^. Complementary proteomic analyses demonstrate significant enrichment of immune, cell-migration, adhesion, and chemotactic pathways in HFpEF RVS tissue, indicating a robust inflammatory component associated with myeloid-cell expansion^65^.

To overcome the limited availability of patient biopsies, we designed the novel murine ‘RV-HFpEF’ model, which develops clinical signs of both HFpEF and RVD. Interestingly, RV-HFpEF does not present with sexual dimorphism as described for the gold-standard two-hit HFpEF model^34^. Given that RV-HFpEF does not rely on genetic manipulation, it enables the combination with cell-specific loss-of-function approaches, e.g. by Cre/lox or Dre/rox systems to study cell-specific molecular mechanisms of RVD in HFpEF. Our model combines classic HFpEF triggers, namely metabolic stress and systemic hypertension, with hypoxia, thus mimicking reduced oxygen levels due to an increase in the alveolar-arterial O_2_ gradient in HFpEF patients^28,29,30,66^.

In our model, RV inflammation, but not RV matrix deposition, was reversed by CSF1R-mediated macrophage depletion, indicating that cells other than those relying on the myeloid receptor CSF1R contribute to matrix deposition in RV-HFpEF. Notwithstanding this, RV hemodynamics improved in response to macrophage depletion independent of collagen content. We speculate that mechanisms beyond collagen deposition, such as direct inflammatory effects impacting cardiomyocyte function, contribute to impaired RV hemodynamics in HFpEF. This notion is supported by the finding of reduced RV neutrophil counts in the absence of macrophages in RV-HFpEF mice. Although neutrophil depletion was not sufficient to rescue HFpEF-associated RVD in our mouse model, neutrophils from HFpEF patients have been shown to release more cytokines compared to controls and LV diastolic dysfunction has previously been linked to neutrophil degranulation and Neutrophil Extracellular Trap (NET) formation in HFpEF^67–69^. Moreover, depletion of resident macrophages via CSF1R inhibition has been shown to reduce neutrophil influx and neutrophil-mediated cytokine release, ultimately mitigating tissue injury^70,71^. Our results from macrophage depletion experiments link macrophage and neutrophil dynamics, indicating cellular crosstalk between these two immune cell populations in HFpEF-associated RVD. The fact that LV dysfunction was not rescued by macrophage depletion may, on the one hand, be attributable to our experimental design, as CSF1R inhibitor treatment was only initiated six weeks after HFpEF induction. On the other hand, this finding is coherent with the fact that HFpEF-LVs rely less on macrophage expansion than HFpEF-RVs. Future studies with preventive macrophage depletion approaches are necessary to study the role of macrophages in LV diastolic dysfunction.

Elevated macrophage counts in HFpEF LVs have been shown to be mediated by the interaction of C-C chemokine receptor type 2 (CCR2) with its ligand monocyte chemoattractant protein-1 (CCL2), which effectively regulate monocyte chemotaxis. Consistently, genetic inhibition of the CCR2/CCL2 axis prevented cardiac macrophage expansion and improved diastolic function in a murine model of angiotensin-II induced HFpEF^51,72^. In our study we found macrophage expansion via recruitment and proliferation of CCR2^+^ myeloid cells in HFpEF RVs at the expense of TIMD4^+^ and LYVE1^+^ resident macrophage subsets. In contrast to recruited macrophage subsets, which promote extracellular matrix deposition after tissue injury^7^, TIMD4^+^ and LYVE1^+^ resident macrophage subsets protect from cardiac tissue remodeling and ventricular dysfunction^73–75^. Our findings are in line with recently published literature on the role of macrophages in LV diastolic dysfunction which demonstrate that resident macrophage subsets are cardioprotective in HFpEF^76,77^, whereas CCR2^+^ monocyte-derived macrophages drive cardiac hypertrophy in HFpEF and are associated with increased extracellular matrix deposition and fibroblast activation, thus promoting myocardial stiffness and impairing myocardial relaxation^7,78^. Of note, modulating CCR2^+^ macrophage subsets by vagal stimulation prevents tissue inflammation and fibrosis in HFpEF LVs^76,79^, highlighting the therapeutic potential of targeting CCR2^+^ macrophages in HFpEF. In contrast to the LV, the role of macrophages in RV remodeling and dysfunction in HFpEF is little understood. While macrophage-associated RV remodeling and RVD have been implicated in several forms of pulmonary arterial hypertension^59,60,80^, our study is the first to demonstrate ventricle-specific differences in macrophage expansion and recruitment in RV-HFpEF, highlighting the need to study disease mechanisms in HFpEF individually for both cardiac chambers.

## Conclusion

Our translational work shows that dysregulated leukocyte counts and myeloid cell dynamics are associated with, and directly contribute to, the pathogenesis of HFpEF-associated RVD. We demonstrate that LV and RV exhibit distinct inflammatory responses, when exposed to the same systemic disease triggers in HFpEF, with HFpEF-RVs being particularly vulnerable to myeloid cell recruitment and sterile inflammation compared to HFpEF-LVs. Immunomodulation of myeloid cells may therefore provide a potential therapeutic approach to prevent RV remodeling and dysfunction in HFpEF.

## Limitation

Our study is limited by the availability of fresh human RV tissue for the validation of leukocyte accumulation in HFpEF patients.

## Acknowledgement

The authors thank all Grune and Kuebler lab members for discussion and insightful comments. The authors thank Daniel Schulze for statistical advice in calculating animal n-numbers and analyzing human data.

## Author contribution

L.J. and C.K. performed and analyzed experiments; interpreted data. L.J. made the figures. A.W., H.K., P.F., P.-L.P., E.A., L.v.O., D.F., J.G. and N.H. performed experiments and collected data. M.M., A.M.C., E.R., V.Z., L.K., D.Z., L.A., K.F., B.P., F.E., V.H. and K.S. provided human specimen and data as well as statistical analyses. W.M.K. and J.G. designed the experiments. S.S., G.G.S., S.v.L., P.S., N.H., W.M.K. and J.G. discussed results and strategy. L.J., C.K. and J.G. wrote the manuscript with input from all authors. J.G. conceived and directed the study.

